# Rapid quantification of tissue perfusion properties with a two-stage look-up table: a simulation study

**DOI:** 10.1101/2021.11.04.467306

**Authors:** Bin Yang

**Affiliations:** Duquesne University, Department of Engineering, Pittsburgh, USA

**Keywords:** spatial frequency domain imaging, hemodynamics, perfusion properties, Monte Carlo simulation, look-up table

## Abstract

Tissue perfusion properties reveal crucial information pertinent to clinical diagnosis and treatment. Multispectral spatial frequency domain imaging (SFDI) is an emerging imaging technique that has been widely used to quantify tissue perfusion properties. However, slow processing speed limits its usefulness in real-time imaging applications. In this study, we present a two-stage look-up table (LUT) approach that accurately and rapidly quantifies optical (absorption and reduced scattering maps) and perfusion (total hemoglobin and oxygen saturation maps) properties using stage-1 and stage-2 LUTs, respectively, based on reflectance images at 660nm and 850nm. The two-stage LUT can be implemented on both CPU and GPU computing platforms. Quantifying tissue perfusion properties using the simulated diffuse reflectance images, we achieved a quantification speed of 266, 174, and 74 frames per second for three image sizes 512×512, 1024×1024, and 2048×2048 pixels, respectively. Quantification of tissue perfusion properties was highly accurate with only 3.5% and 2.5% error for total hemoglobin and oxygen saturation quantification, respectively. The two-stage LUT has the potential to be adopted in existing SFDI applications to enable real-time imaging capability of tissue hemodynamics.

## 1. Introduction

Sufficient oxygen supply in tissue is critical to maintaining its normal physiological functions [1, 2]. Local oxygen levels in tissue provide valuable insights into its health states, such as metabolic levels [3]. Oxygen is transported via binding to hemoglobin, the main protein of red blood cells, to different parts of the body, and is released to tissue through perfusion. Depending on the presence of binding oxygen molecules, hemoglobin can exist in two forms, oxygenated and deoxygenated hemoglobin, both of which exhibit distinct absorption properties in visible and near-infrared (NIR) ranges. Based on the differential absorption of hemoglobin, multispectral imaging was developed to quantify tissue perfusion properties, such as oxygen saturation[4]. However, the quantification accuracy was limited due to its inability to decouple tissue absorption from tissue scattering. Spatial frequency domain imaging (SFDI), on the other hand, independently evaluates tissue absorption and scattering properties using 2D structured illuminations [5-7]. Multispectral SDFI has been developed to map perfusion and hemodynamic properties of the tissue with an improved accuracy [8, 9]. Such quantification requires imaging at multiple wavelengths (two wavelengths at least), typically in the red and near-infrared (NIR) range [10, 11].Typically, at each wavelength, a set of 3 phase-shifted patterns are projected to the tissue surface and the corresponding reflectance images are acquired to determine the tissue’s optical and perfusion properties [6].

The overall process, however, is slow and limits the temporal resolution. In the past few years, significant efforts have been devoted to improving the imaging and quantification speed in SFDI imaging. Single snapshot SFDI reduced the required images from 3 to 1 [12-14], and look-up table (LUT) based quantification significantly improved the quantification speed of optical property of the tissue [15]. For perfusion property quantification, Beer-Lambert Law is the standard approach, which could be slow for large images. To improve the speed for perfusion property quantification, artificial intelligence was employed to train the predictive model based on a large set of training images [16, 17].

Despite recent advancements towards real-time mapping of tissue perfusion and hemodynamics, current methods are still limited by relatively slow processing speed, low image resolution, and the need for powerful computational hardware (e.g., multi-GPU setup). This study aimed to develop a highly efficient LUT-based computational framework towards high-resolution real-time mapping of tissue perfusion and hemodynamics that can be implemented on cost-effective hardware platforms.

## 2. Methods

LUT techniques have been widely used in SFDI to determine optical properties without performing time-consuming fitting procedures [18]. However, LUT techniques were rarely used to quantify tissue perfusion properties in conjunction with SFDI imaging. In this study, we developed a highly efficient two-stage LUT technique consisting of two connected LUTs. The first and second stage LUTs rapidly quantify tissue optical and perfusion properties, respectively. Fig. 1 summarizes the overall workflow of the two-stage LUT technique.

**Fig. 1.**
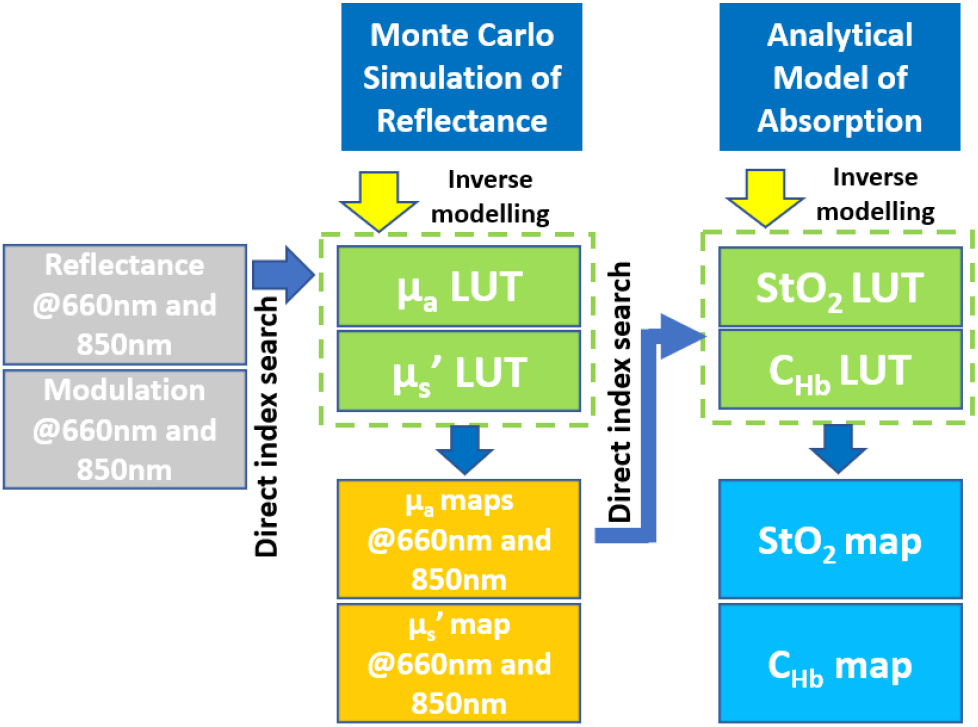
Overview of two-stage LUT. Monte Carlo simulation of reflectance and analytical model of blood absorption are inverted to generate LUTs for optical properties mapping (stage-1) and perfusion mapping (stage-2), respectively. SFDI measurements at 660nm and 850nm are used as inputs to quantify absorption and reduced scattering coefficients. The absorption coefficients are later used as search indices to quantify oxygen saturation and total hemoglobin concentration.

The stage-1 LUT was generated by inverse modeling diffuse reflectance obtained by Monte Carlo simulation in the spatial frequency domain, and the stage-2 LUT was generated by inverse modeling of an analytical model of tissue absorption. Quantification of tissue perfusion properties was based on SFDI imaging at 660nm and 850nm. These two wavelengths located on the opposite side of the isosbestic wavelength, around 800nm, have been previously demonstrated to quantify blood perfusion properties effectively [16]. With the two-stage LUT, reflectance and modulation measurements at both 660nm and 850nm were first used as search indices to determine absorption and reduced scattering coefficient using the stage-1 LUT. The absorption coefficients at 660nm and 850nm were subsequently used as searching indices to efficiently determine oxygen saturation and blood concentration using the stage-2 LUT. In the rest of the section, detailed procedures of generating and evaluating the performance of two-stage LUT for quantifying perfusion properties will be presented.

### 2.1 Stage-1 LUT for quantifying optical properties

Monte Carlo (MC) simulation has been widely used for simulating light transport in turbid media. The simulation results are often used to generate LUT tables for quantifying optical properties through inverse modeling [19, 20]. Traditionally, MC simulation was implemented on CPU [21], and is considered computationally intensive and slow [22]. Due to its highly parallel nature of execution, graphic computing unit (GPU) based MC simulation programs have recently been developed, which significantly improve the simulation efficiency [23]. This study adopted the Monte Carlo eXtreme (MCX) simulation package developed by Fang and Boas to simulate reflectance in the spatial frequency domain [24].

We simulated reflectance images under structured illumination with a spatial frequency of 0.125mm^-1^ on large virtual phantoms (9.6cm x 9.6cm x 5cm) with a grid size of 0.1mm in x, y, and z directions. The sample thickness of 5cm was chosen to satisfy the requirement for infinite geometry in MC simulations [18]. To reflect the general properties of biological tissue, the virtual phantom was assumed to have an anisotropy g=0.9, and a refractive index n=1.37 [25]. A total of 208 pairs of biologically relevant optical properties were simulated with 16 absorption coefficient (μ_a_) values from 0.001mm^-1^ to 1mm^-1^, and 13 reduced scattering coefficient values (μ_s_’) from 0.5mm^-1^ to 3mm^-1^. For each sample, diffuse reflectance images were simulated under 3 phase-stepped (0°, 120°, and 240°) structured illuminations. Thus, a total of 624 MC simulations were performed, and the corresponding reflectance images were saved. To reduce the noises in simulation, a total of 200 million photons was launched in each simulation.

Following the simulation, a sub-region (red-dash square in Fig. 2A) of each reflectance image was isolated to reduce the signal fall-off toward the edge of the image. The resultant diffuse reflectance image was then averaged along the pattern direction (yellow arrows in Fig. 2A) to further improve the signal quality. Three intensity profiles, *I*_*1*_, *I*_*2*_ and *I*_*3*_, were obtained for three phase-stepped illuminations (Fig. 2B). The demodulation process, shown in Eq. 1 and 2 was then performed to extract DC (*I*_*DC*_) and AC (*I*_*AC*_) signals (Fig. 2c).

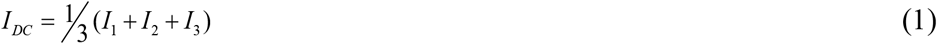

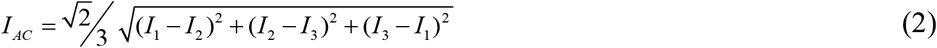

**Fig. 2.**
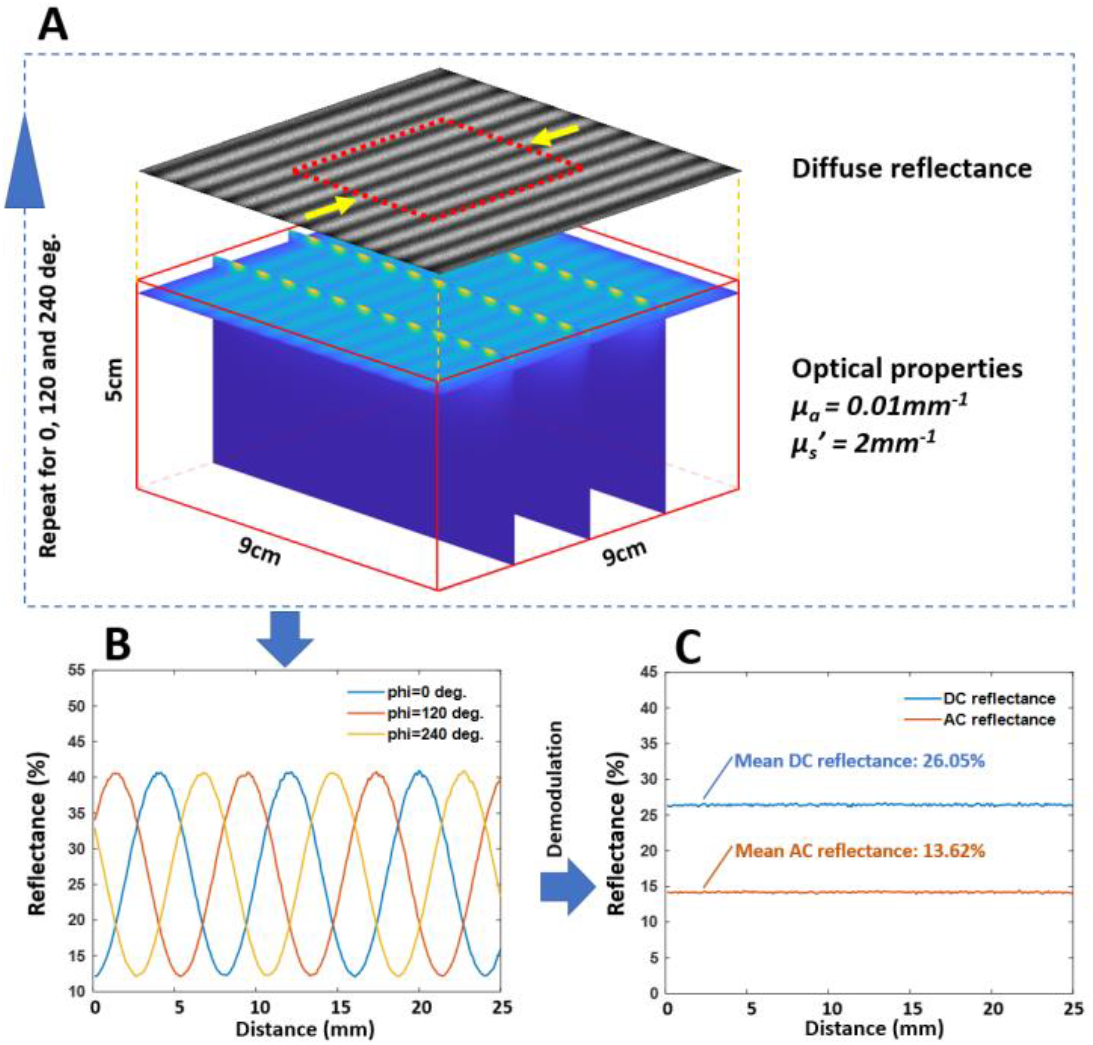
Monte Carlo simulation workflow. A.Schematic of Monte Carlo simulation over a large volume. For each setting, the simulation was performed with phase shifts of 0, 120, and 240 degrees. The central region (red dash box) of the diffuse reflectance image (gray-scale image) was averaged along the direction of yellow arrows to obtain reflectance plots. B. Reflectance plots at three phase shifts. C. Demodulated AC and DC intensity plots.

*I*_*DC*_ and *I*_*AC*_ signals were further converted to reflectance (R) and modulation (M) with Eq. 3 and 4, respectively.

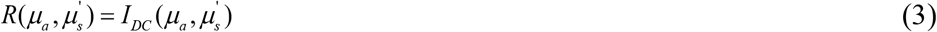

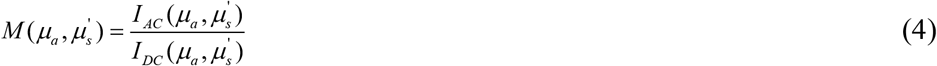

Maps of 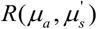 and 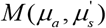 were converted to LUTs, 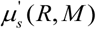 and 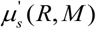, using a coordinate mapping technique similar to the one described elsewhere [15]. Briefly, after obtaining 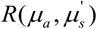 and 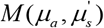 from the simulation, a 2D interpolation was performed to generate another set of R and M maps with a finer and uniform grid size of μ_a_ and μ_s_’. Data points in the newly generated 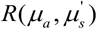 and 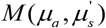 maps were directly mapped to generate two new maps *µ*_*a*_ (*R, M*) and 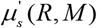. A scattered data interpolation was used to fill the gaps in *µ*_*a*_ (*R, M*) and 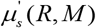 maps. To facilitate a rapid search for optical properties, during the LUT generation, reflectance and modulation were up-scaled by a factor of 1000, and rounded to the nearest integer. This operation allowed for direct index searching based on reflectance measurements.

### 2.2 Stage-2 LUT for quantifying perfusion properties

We performed forward modeling of tissue absorption at two wavelengths, 660nm and 850nm. Hemoglobin concentration, *CHb*, was varied from 0.075 g/L to 22.5 g/L with an increment of 0.075g/L, which corresponds to 0.05% and 15% of the total hemoglobin concentration of 150 g/L in the whole blood. This *CHb* range reflects the majority types of organs except for the kidney and the liver [25]. *CHb* in those two organs is well above 30% [25]. Oxygen saturation, *StO*_*2*_, was varied from 50% to 100% with an increment of 0.25%. A hemoglobin molecular weight of 64500 g/mole was used in the simulation [26]. The absorption coefficient at both wavelengths can be analytically described by Eq. 5 and 6, respectively, where 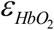 and *ε* _*Hb*_ are molar extinction coefficients of oxygenated and deoxygenated hemoglobin. With a similar technique described in the previous section, maps of *µ*_*a* _ 660_ (*StO*2, *CHb*) and *µ*_*a* _ 850_ (*StO*2, *CHb*) were converted to maps of *StO*2(*µ*_*a* _ 660_, *µ*_*a* _ 850_) and *CHb*(*µ*_*a* _ 660_, *µ*_*a* _ 850_), and a scaling factor of 5000 was used.

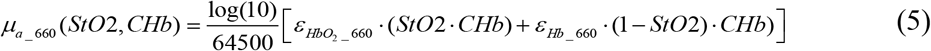

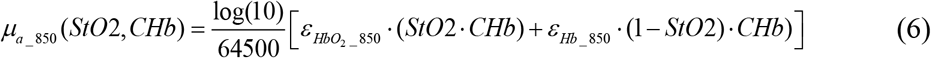

### 2.3 Quantification speed test

Speed of quantifying optical and perfusion properties using the two-stage LUT was evaluated based on a set of simulated diffuse reflectance images (Section 2.3.1 and 2.3.2). The performance of stage-2 LUT was compared to the Beer-Lambert Law based quantification (Section 2.3.3). All quantification methods were implemented on both CPU and GPU to fully evaluate their efficiency. GPU acceleration was achieved by simply converting a regular MATLAB array into a GPU-compatible array with the MATLAB function `gpuArray()`. The testing platform consists of an Intel CPU (i9 10900K), 32 GB of memory, and a Nvidia GPU (RTX 3080). A speed test script was written in MATLAB, and only the built-in MATLAB functions were used. We performed a speed test on three sets of simulated images with resolutions 512×512 pixels, 1024×1024 pixels, and 2048×2048 pixels. Each set of images consisted of 50 images. Three image sizes were later referred to as small, medium, and large sizes, respectively.

#### 2.3.1 Optical property mapping using stage-1 LUT

Reflectance and modulation images obtained from the demodulation were first multiplied by a scaling factor of 1000, and rounded to the nearest integer. MATLAB function `sub2ind()` was used to convert 2D maps of scaled reflectance and modulation into linear search indices for stage-1 LUT to quantify optical properties, as shown in Fig. 3A and 3B.

**Fig. 3.**
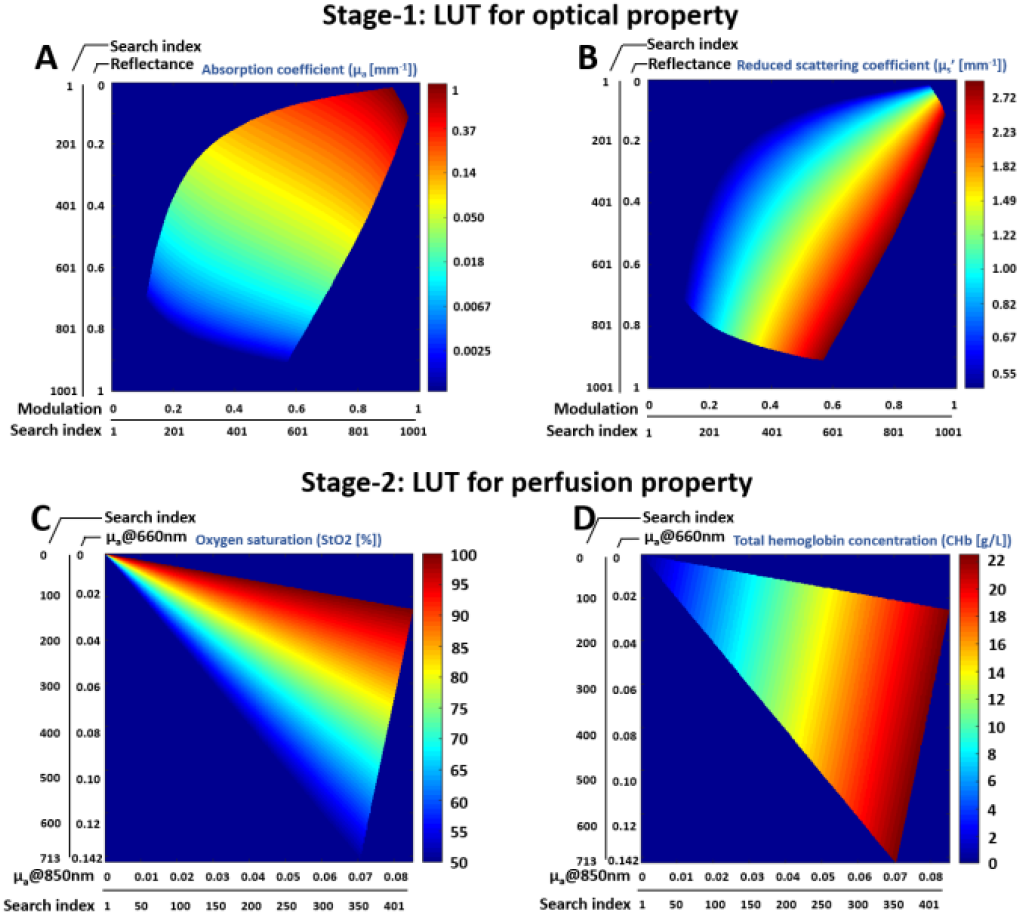
Two-stage LUTs for stage-1 optical (A and B) and stage-2 perfusion (C and D) properties mapping. Reflectance and modulation in stage-1 LUT, and absorption coefficients at 660nm and 850nm in stage-2 LUT were converted into search indices.

#### 2.3.2 Perfusion property mapping using stage-2 LUT

Following optical property quantification, absorption coefficient maps at 660nm and 850nm were multiplied by a scaling factor of 5000, and similarly, the integer portion of the result was retained. MATLAB function `sub2ind()` was used to convert 2D maps of scaled absorption coefficient maps into linear search indices for stage-2 LUT to quantify perfusion properties, as shown in Fig. 3C and 3D.

#### 2.3.3 Perfusion property mapping using Beer-Lambert Law

Eq. 5 and 6 were solved for concentration of oxygenated hemoglobin (= *StO*2. *CHb*) and deoxygenated hemoglobin (= (1− *StO*2). *CHb*), based on which oxygen saturation and total hemoglobin concentration were determined. To improve the computation efficiency, 2D absorption maps were first converted to 1D array to take the advantage of high efficiency of vectorized computation in MATLAB.

### 2.4 in-silico validation

We examined the quantification accuracy of the perfusion parameters through an *in-silico* validation procedure. We generated five sets of digital phantoms with total hemoglobin concentrations varying from 2.25g/L to 20.25g/L with an increment of 4.5g/L. At each hemoglobin concentration, five phantoms were created with the oxygen saturation levels varying from 50% to 100% with an increment of 10%. Thus, a total of 30 phantoms were generated. Absorption coefficients at 660nm and 850nm were calculated using Eq. (5) and (6) for all phantoms. The absorption coefficients were later used to quantify hemoglobin concentration and oxygen saturation using the stage-2 LUT. Quantification errors were calculated based on ground truth values.

## 3. Results

### 3.1 Two-stage LUT

Figure 3 exhibits a two-stage LUT for optical and perfusion property mapping. The axes of each LUT are characterized by actual values and their corresponding search indices. In both stages, the corresponding search indices are used to determine the optical and perfusion properties.

### 3.2 Quantification speed test

Table 1 summarizes the processing time for quantifying optical properties and perfusion properties using both GPU- and CPU-based computation. It is clear that with the GPU acceleration, the processing time of three types of quantification have been reduced. Note that the LUT access time (processing time in the parentheses) accounts for a smaller portion of the over processing time in the GPU-based processing than that in CPU-based processing.

**Table 1.** Processing time for optical and perfusion property quantification with GPU and CPU.

|  |  | GPU-based |  |  | CPU-based |  |  |
| --- | --- | --- | --- | --- | --- | --- | --- |
| Image size (pixels x pixels) |  | 512(S) | 1024(M) | 2048(L) | 512(S) | 1024(M) | 2048(L) |
| Processing time (ms) | I. $\mu_a$ and $\mu_s'$ mapping at 660nm and 850nm with stage-1 LUT | 2.36<br>(0.87)* | 3.39<br>(1.04) | 6.94<br>(2.47) | 10.64<br>(8.19) | 41.72<br>(31.05) | 170.34<br>(128.84) |
| | II. $StO_2$ and CHb mapping with stage-2 LUT | 1.41<br>(0.19) | 2.34<br>(0.22) | 6.49<br>(0.36) | 4.87<br>(2.69) | 20.52<br>(10.84) | 86.61<br>(45.96) |
| | III. $StO_2$ and CHb mapping using Beer-Lambert Law | 9.18 | 24.72 | 93.33 | 9.48 | 37.91 | 165.81 |

\*GPU access time.

While the quantifications for the perfusion property with both stage-2 LUT and Beer-Lambert law were accelerated with a GPU, stage-2 LUT clearly showed a much higher efficiency. The stage-2 LUT is approximately 5.5 times and 13.5 times faster for small and large sized images, respectively. Without GPU acceleration, stage-2 LUT still showed about 90% higher efficiency than the Beer-Lambert Law method.

It is worth noting that a large image size imposes a much higher penalty on CPU-based processing than GPU-based processing. The CPU-based processing time is proportional (1x, 4x, and 16x) to the image sizes for all three types of quantifications. On the other hand, the GPU-based processing time is less affected by the image sizes, especially for the LUT-based quantifications

Converting the processing time to processing speed (frames per second/FPS) revealed GPU-based two-stage LUT (sum of processing time I and II) was capable of achieving a processing speed of 266, 174, and 74 FPS for small, median, and large sized images, respectively, which is 2.1, 3.9, and 6.5 times faster than that of the Beer-Lambert Law based processing (sum of processing time I and III) (Fig. 4A). On the other hand, CPU-based quantification was slower, but it still achieved approximately 65, 16, and 4 FPS with two-stage LUT for three image sizes, which is about 30% faster than Beer-Lambert law-based processing (Fig. 4B). Note that the processing time of optical properties includes both absorption and reduced scattering coefficients. If tissue scattering property is not of interest, excluding scattering quantification could further improve the quantification speed.

**Fig. 4.**
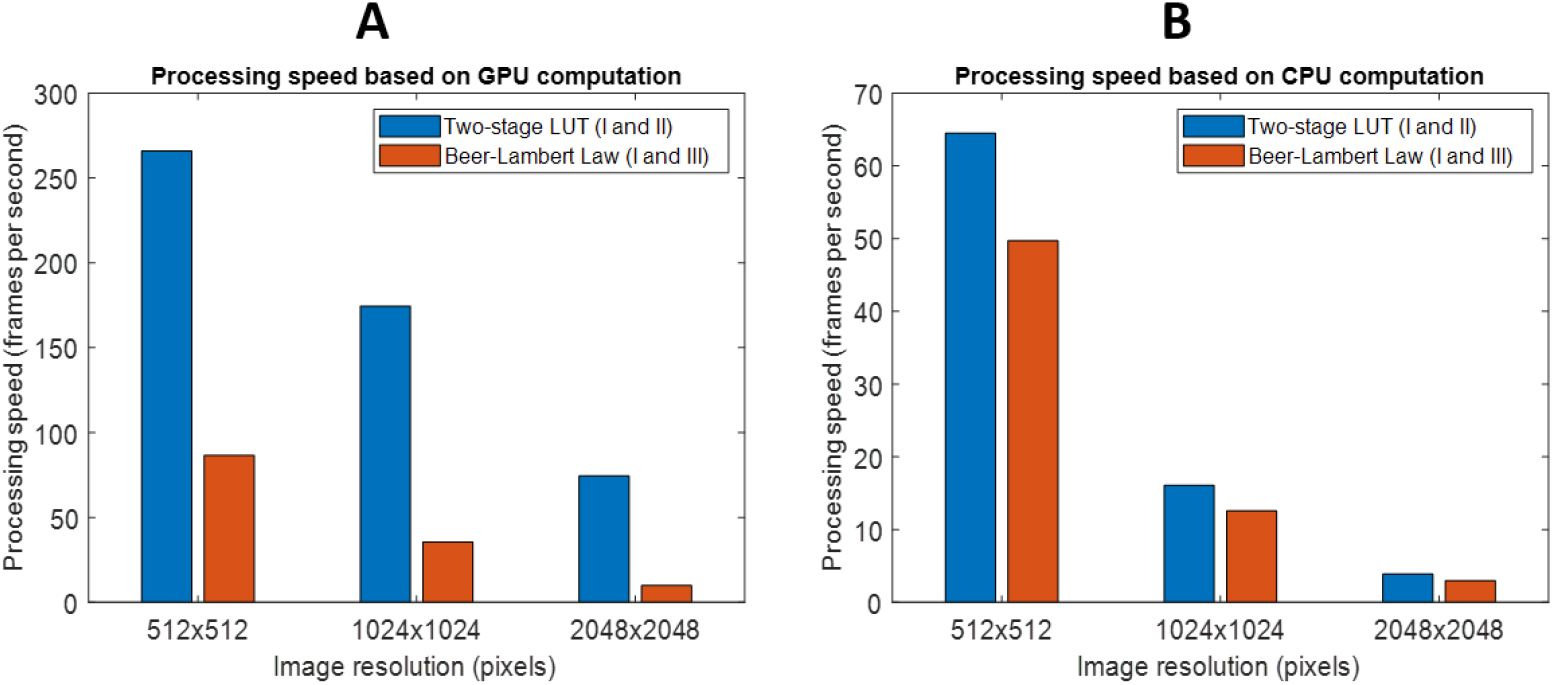
A. Quantification frame rate of optical and perfusion properties of three image sizes with a GPU and a CPU. B. Processing time increases as the image size increases for both GPU- and CPU-based processing. However, the GPU-based is less affected.

### 3.3 in-silico validation

The stage-2 LUT exhibited a high accuracy of quantifying perfusion properties as shown in Fig.5. The absolute quantification error is less than 3.5% for total hemoglobin concentration and less than 2.5% for oxygen saturation over a wide range of perfusion properties. While the overall error is small, Fig. 5 suggests that a lower total hemoglobin concentration may correlate with a higher error.

**Fig. 5.**
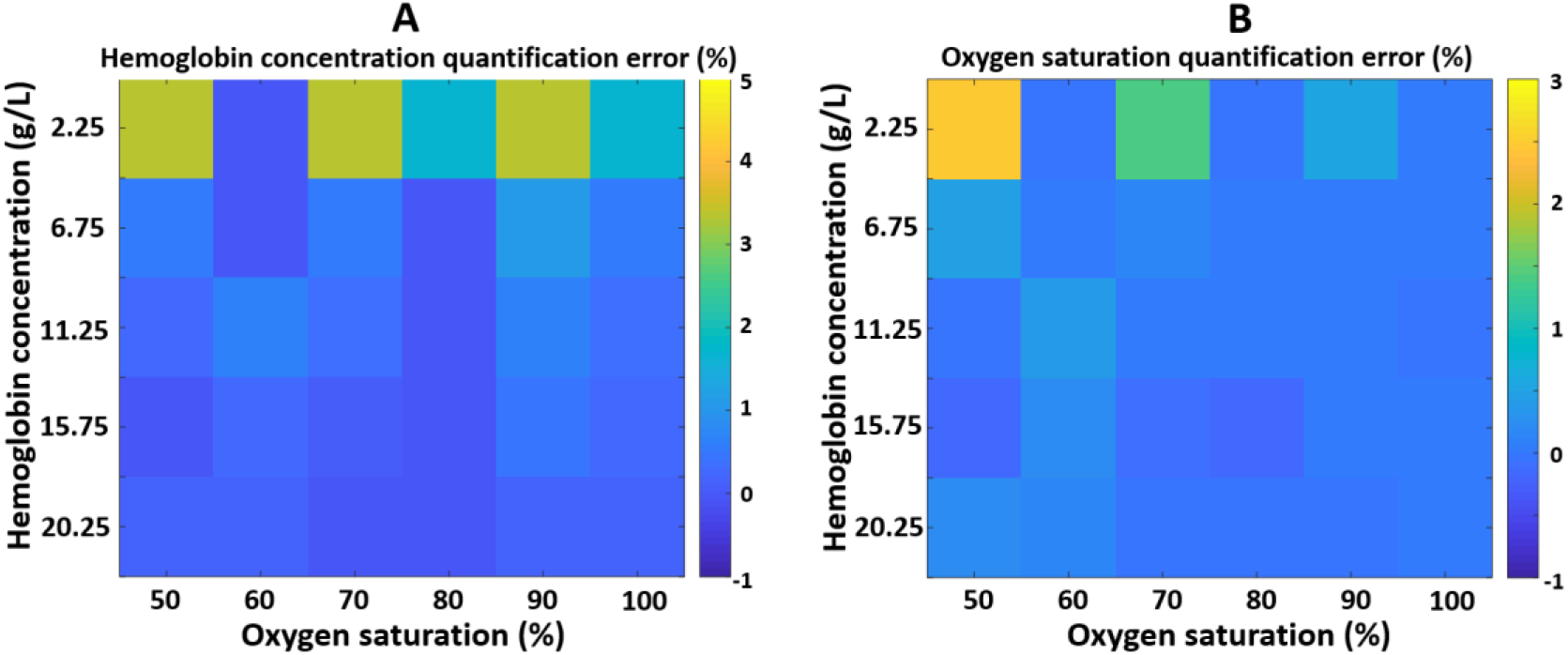
Quantification accuracy analysis of hemoglobin concentration and oxygen saturation with stage-2 LUT.

## 4. Discussion

In this study, we developed and demonstrated a highly efficient and accurate two-stage LUT method that quantifies the tissue’s optical and perfusion properties with SFDI. High efficiency was achieved through direct index searching of two LUTs, which eliminated time-consuming fitting operations to determine optical and subsequently perfusion properties. GPU-based implementation significantly improved the searching speed through parallelization. At an image size of 2048×2048 pixels, we achieved a quantification speed of 74 FPS for both optical and perfusion properties.

In recent years, methods for direct quantification of optical and perfusion properties using artificial intelligence have been reported to improve the processing speed towards real-time applications. Zhao et al. reported a processing time of 94.4ms (excluding time for image import and result display) for quantifying perfusion properties for images with a size of 540×720 pixels using a deep residual network [17]. Using adversarial deep learning, Chen and Durr reported that approximately 40ms was need to infer optical and perfusion properties for images with a size of 512×512 pixels [16]. It is worth noting that both studies utilized two wavelengths at around 660nm and 850nm, which are the same as the wavelengths used in this study.

A two-stage LUT method is highly suitable for real-time applications. With GPU acceleration, it achieved 266 FPS and 74 FPS for small and large sized images. The short processing time could provide ample room to accommodate more sophisticated algorithms while still maintaining real-time performance. When the two-stage LUT is coupled with a dual-sensor SFDI imaging system, reflectance images at 660nm and 850nm could be acquired simultaneously and processed rapidly, which would enable time-sensitive imaging applications, such as real-time tissue viability monitoring.

Two-stage LUTs outperformed the Beer-Lamber Law based quantification regardless of the computational platform. Even with GPU acceleration, Beer-Lamber Law method only achieved 35.5 FPS for median size images. Any additional processing would slow it down to sub-30 FPS, which is not ideal for real-time applications.

While CPU-based implementation of two-stage LUT is not as fast, it does not require a dedicated GPU and offer a few advantages. First, the two-stage LUT is still suitable for offline applications where processing time is not a critical factor, and it is about 30% more efficient than Beer-Lamber Law approach. Second, the two-stage LUT offered approximately a processing speed of 65 FPS for small sized images, which still could enable real-time applications when the image resolution is not a concern. It is worth noting that in real life applications, acquiring and transferring images to memory, and displaying the processed results take additional time. To preserve the high performance of two-stage LUT, multi-threaded architecture could be adopted. Each thread runs independently and is only responsible for its own task, such as image transfer, quantification, or result display.

The two-stage LUT quantification method presented in this study was intended to serve as a general framework to map the optical and perfusion properties of the tissue. Implementing this general framework requires additional considerations. Firstly, the optical property LUT was generated using MC simulation on homogenous phantoms. While MC simulation is highly accurate, the LUT is not imaging system-specific. When applying this LUT to quantify optical properties of the sample imaged with a specific SFDI system, additional calibration and scaling of the LUT are likely needed to match the characteristics of the imaging system and to achieve a high quantification accuracy. Secondly, the sample’s 3D surface shape affects the diffuse reflectance, and thus the quantification, which was not accounted for in our method. Multiple techniques have been reported to correct for surface height, and these methods could be adopted to improve quantification accuracy [13, 27]. Finally, the two-stage LUT was based on two wavelengths that are sensitive to oxygenated and deoxygenated hemoglobin. If the tissue to be measured presents additional major chromophores, such as melanin, this method may not yield a high quantification accuracy. Higher-dimensional (e.g., three dimensional) LUT could be generated to account for additional chromophores.

## 5. Conclusion

In this work, we presented a highly efficient method, two-stage LUT, that quantifies the tissue’s optical and perfusion properties in SFDI imaging. Both GPU- and CPU-based implementations demonstrated a high efficiency. This method can potentially be adapted and incorporated into existing SFDI imaging applications to improve data processing efficiency. In future work, we will implement this two-stage LUT in a dual-wavelength SFDI imaging system to monitor tissue perfusion and hemodynamics in real-time.

## Acknowledgments

This research work was supported by the Department of Engineering and Faculty Startup Funds provided by the Provost Office at Duquesne University. The author also would like to thank William Miller at Duquesne University for editing and proofreading the manuscript.

## Data availability

The data and code that support the findings of this study are available from the corresponding author upon reasonable request.

